# Unveiling Fine-scale Spatial Structures and Amplifying Gene Expression Signals in Ultra-Large ST slices with HERGAST

**DOI:** 10.1101/2024.08.09.607422

**Authors:** Yuqiao Gong, Xin Yuan, Qiong Jiao, Zhangsheng Yu

**Author notes:** These authors contributed equally: Yuqiao Gong, Xin Yuan.

## Abstract

We propose HERGAST, a system for spatial structure identification and signal amplification in ultra-large-scale and ultra-high-resolution spatial transcriptomics data. To handle ultra-large ST data, we consider the divide and conquer strategy and devise a Divide-Iterate-Conque framework specially for spatial transcriptomics data analysis, which can also be adopted by other computational methods for extending to ultra-large-scale ST data analysis. To tackle the potential oversmoothing problem arising from data splitting, we construct a heterogeneous graph network to incorporate both local and global spatial relationships. In simulation, HERGAST consistently outperformed other methods across all settings with more than 10% average gaining. In real-world data, HERGAST’s high-precision spatial clustering enabled finding SPP1+ macrophages intermingled in tumors in colorectal cancer, while the enhanced gene expression signal enabled discovering unique spatial expression pattern of key genes in breast cancer.

## Introduction

Spatial transcriptomics (ST) technologies have transformed our understanding of gene expression organization within tissues, providing valuable insights into cellular heterogeneity and tissue architecture. Traditional methods like 10X Genomics’ Visium ^1^ have enabled gene expression mapping across tissue sections, offering crucial spatial context. Recently, 10X Genomics introduced groundbreaking advancements with their ultra-large-scale Visium HD ^2^ (commonly about 500K spots, Table S1) and ultra-high-resolution Xenium ^3^ (reaching subcellular resolution) platforms. Compared to previous technologies, these advancements represent an exponential increase in data scale, posing significant analytical challenges. Despite the progress made through various computational approaches in analyzing spatial transcriptomics data, the advent of these new ultra-large-scale and ultra-high-resolution technologies presents substantial challenges. Existing methods, including statistical models like BayesSpace^4^ and SpatialPCA^5^, and deep learning approaches utilizing graph neural networks (GNNs) such as SpaGCN ^6^, SEDR ^7^, conST ^8^, STAGATE ^9^, and GraphST ^10^, have demonstrated impressive performance on previous datasets. However, these methods are no longer scalable or applicable to the new datasets generated by Visium HD or Xenium (Figure S1a-b). Additionally, the increased resolution and data scale dilute biological signals, with highly expressed genes overshadowing signals from lowly expressed ones in Xenium data ^3^. Enhancing or denoising the biological signal in such large-scale, high-resolution datasets is also a pressing issue that must be addressed.

## Results

### HERGAST model overview

The predominant challenges lie in the unsuitability of current algorithms tailored for spatial transcriptomics due to the overwhelming data volume. Simpler techniques like PCA, focusing solely on gene expression similarity, inadequately harness spatial information, thus limiting the comprehensive utilization of spatial transcriptomics data. Drawing inspiration from the divide and conquer strategy prominent in computer science and mathematics^11^, we devise a “Divide-Iterate-Conquer” (DIC) framework meticulously crafted for the analysis of ultra-large-scale spatial transcriptomics data. This involves splitting the slice into manageable patches, iteratively training on these patches, and then inferring on the whole slice (Figure 1a, see Methods). This strategy ensures scalability across arbitrarily large datasets. Additionally, we utilize a heterogeneous graph network that integrates both spatial proximity and gene expression similarity (Figure 1b). By considering spatial proximity relation, spots can be aware of the surrounding local niches and take advantage of the spatial information. By considering gene expression similarity relation, spots can attend to distant spots with similar expression profiles, mitigating over-smoothing issue may be caused by data splitting. Through a cross-attention mechanism, our model adaptively learns attention weights for different relationships, allowing it to incorporate both local and global spatial relationships. By generating low-dimensional embeddings that incorporate both gene expression similarity and spatial proximity, our model enables delineating fine spatial structure. Furthermore, the reconstructed gene expression profiles, informed by these spatial distribution relationships, enhance critical spatial patterns and amplify biologically significant signals that may be subtle in the original data (Figure 1c). Our heterogeneous graph network model, within the specially designed DIC framework, is named HERGAST (High-resolution Enhanced Relational Graph Attention Network for Spatial Transcriptomics).

**Figure 1.**
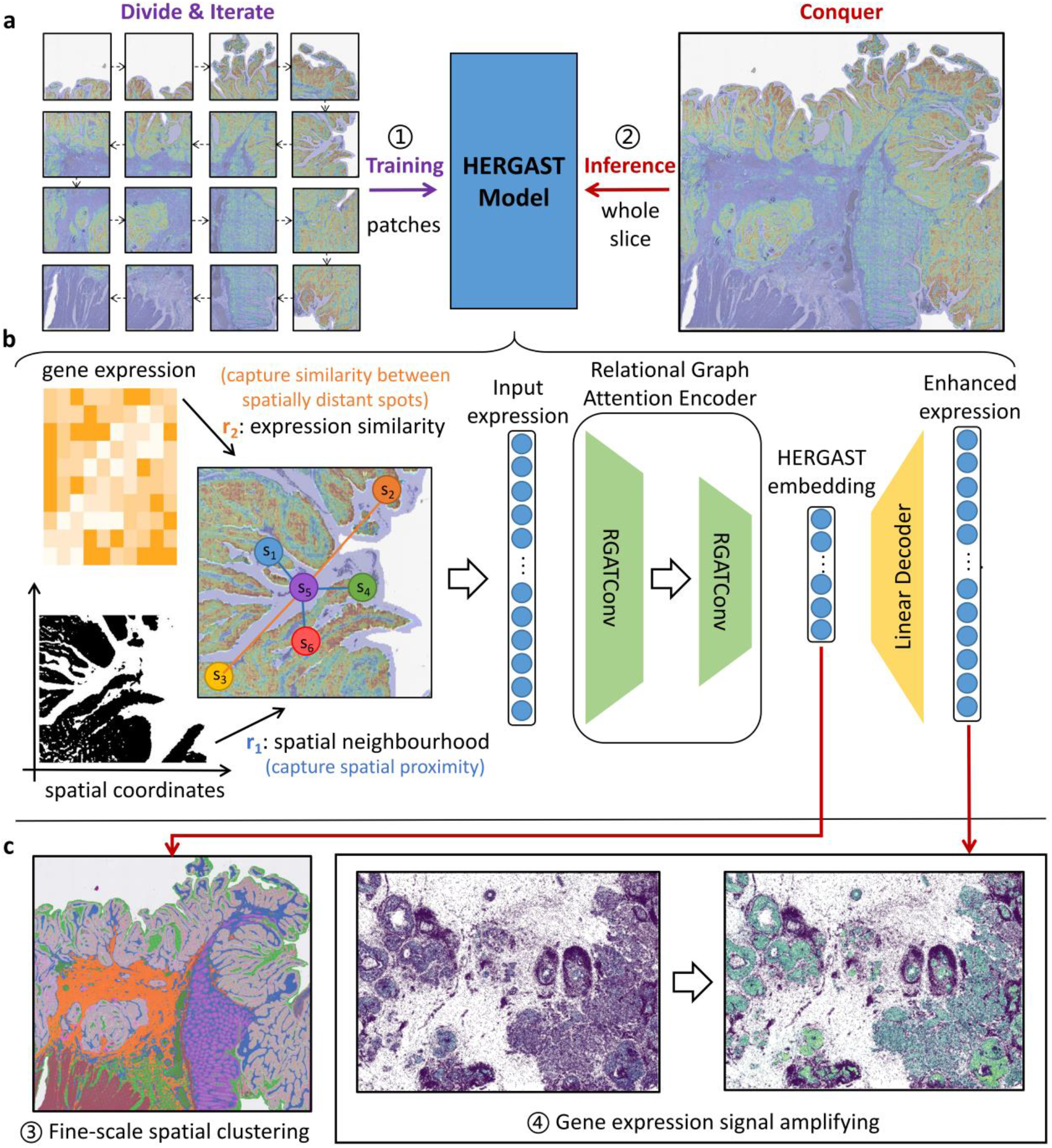
Overview of the HERGAST model. **a**, We divide the spatial transcriptomics data into patches and iteratively train HERGAST on them, inference is then conducted on the entire slice. **b**, HERGAST utilizes spatial information and gene expressions to learn low-dimensional latent embeddings. Each spot’s profile is transformed into a latent embedding by an encoder and reconstructed using a linear decoder. **c**, By considering gene expression similarity and spatial proximity, HERGAST generates low-dimensional embeddings that enable fine-scale spatial clustering. The reconstructed expression profile serves as amplified gene expression signal.

Previous graph network-based methods could theoretically leverage the DIC strategy for ultra-large-scale spatial transcriptomics datasets. However, evaluations reveal significant challenges faced by methods like GraphST, SEDR, and conST, which rely on graph convolutional networks (GCNs). These methods require storing an adjacency matrix for spot relationships and performing matrix multiplications during convolution operations^7,8^,10. Even if we split the data into patches during training, computational and memory requirements at the inference stage make these methods impractical for benefiting from the DIC strategy. To illustrate this point, we conducted tests while imposing a maximum memory usage constraint of 512G and examined the corresponding maximum number of supported spots for each method when adopting the DIC strategy. GraphST, SEDR, and conST demonstrate limited scalability that falls short of practical usage (Figure S1a). Among the existing methods, only STAGATE, which employs a “message passing” mechanism similar to our graph attention network, shows promise for scaling with the DIC strategy ^9^ (Figure S1c). We applied the DIC strategy to STAGATE, subsequently including it in our comparative analysis as STAGATE(DIC). Additionally, we incorporated PCA (implemented by Scanpy package^12^), a straightforward dimensionality reduction method only considering expression similarity, as a baseline for comparison.

### HERGAST consistently outperformed other methods in simulation

We first conducted a simulation to compare HERGAST with PCA and STAGATE(DIC). Spatial tissue structures were manually generated, composed of different cell clusters. As the number of spots increased, the complexity of the spatial tissue structure and the variety of cell clusters also increased, reflecting the advancements in spatial transcriptomics (Figure 2a, Figure S2). Gene expressions for each cell cluster were randomly sampled from the integrated Human Lung Cell Atlas (HLCA), which includes cell annotations ^13^ (see Methods). We used the ARI (Adjusted Rand Index) to evaluate the performance of HERGAST and the other methods. HERGAST (average ARI=0.609) consistently outperformed both STAGATE(DIC) (average ARI=0.487) and PCA (average ARI=0.375) across all simulations, maintaining a relatively high ARI above 0.6 even as the number of spots increased (Figure 2b). This indicates robust performance in large-scale data scenarios. Conversely, PCA’s performance declined sharply with intricate patterns and larger data scales, showing that methods relying solely on gene expression without considering spatial relationships are inadequate for large-scale spatial transcriptomics data. Although STAGATE(DIC)’s ARI metric did not deteriorate as severely as PCA’s, over-smoothing of spatial domains was observed, indicating its inability to capture heterogeneity in large-scale, high-resolution spatial transcriptomics data (Figure 2b, Figure S2).

**Figure 2.**
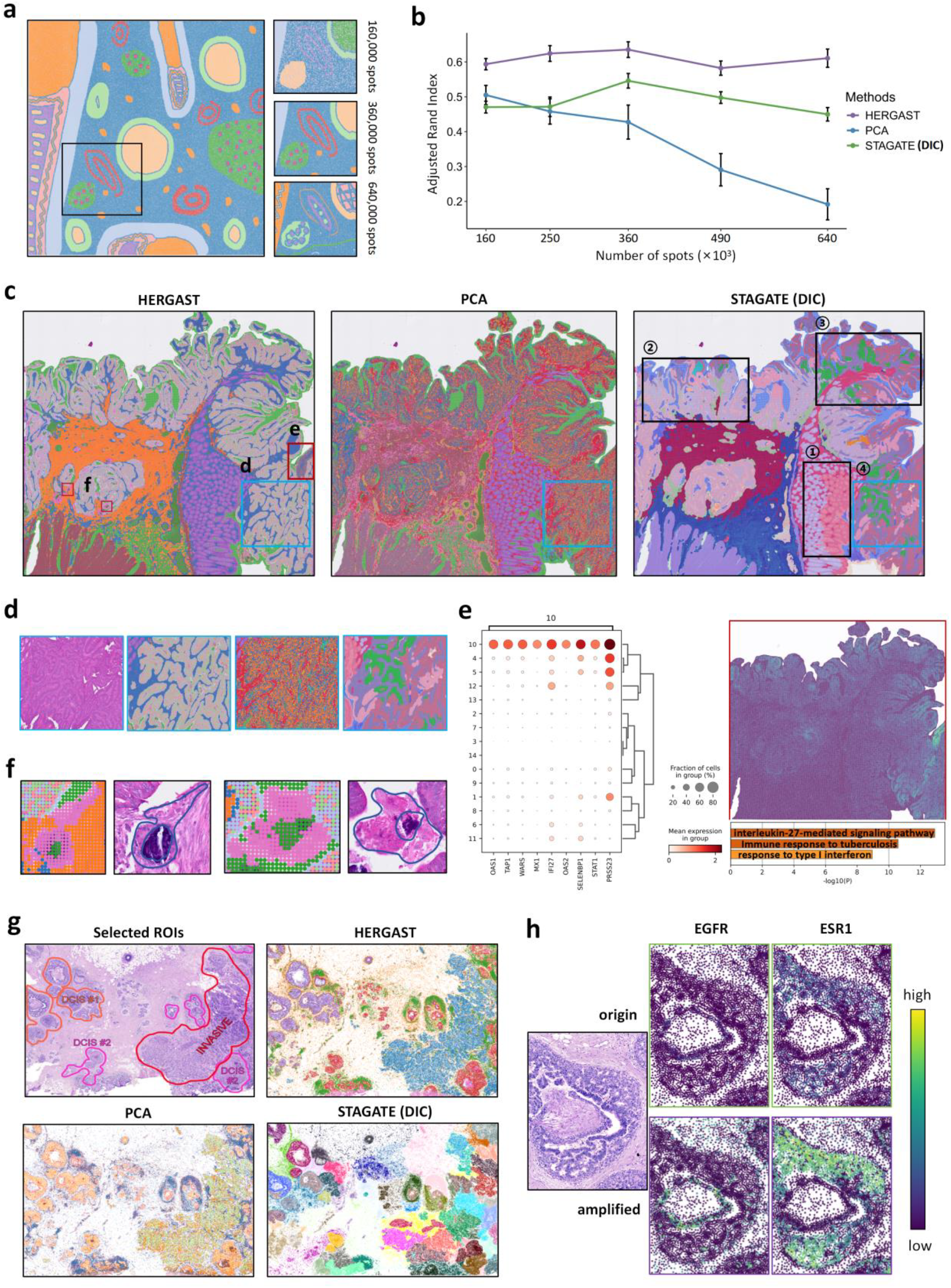
Benchmarking and application of HERGAST. **a**, An illustration of the simulated spatial transcriptome data. The left panel depicts the ground truth spatial pattern in a setting with 360,000 spots. The right panel demonstrates the increasingly complex spatial pattern as the number of spots increases, exemplified by the indicated area in the left panel. **b**, Performance of spatial clustering of different methods across varying conditions. *x* ™axis represents the number of spots in the corresponding simulation setting and y axis represents the ARI score. We ran 10 independent replications for each setting, the data points represent mean value and the error bars represent the standard error calculated across these replications. Visualization results of last replication in each setting can be found in Figure S2. Results for all 10 replications can be found in Table S2. **c**, Spatial clustering results of different methods in Visium HD human colorectal cancer slice. **d**, A zoomed-in view of a tumor area with hematoxylin and eosin (H&E) stained image (first panel), HERGAST result (second panel), PCA result (third panel) and STAGATE(DIC) result (last panel). **e**, Left: dot plot of the highly expressed genes in HERGAST’s unique cluster indicated as box e in Figure 2c. Right top: spatial expression of the combined metagene. Right bottom: corresponding significantly enriched biological pathways determined with the Metascape web tool. **f**, Zoom-in view of two example of the spatial cluster of phagocytes uniquely identified by HERGAST (indicated as box f in Figure 2c) and the corresponding H&E image. More example can be found in Figure S4. **g**, H&E image and spatial clustering results of different methods in Xenium human breast cancer slice. **h**, Spatial expression of EGFR and ESR1 of the original (right top) and amplified (right bottom) expression of a selected region (left). More examples can be found in Figure S6.

### HERGAST enable fine-scale tumor microenvironment mapping in colorectal cancer

We then compared the spatial clustering results of HERGAST, PCA, and STAGATE(DIC) on a Visium HD slice from a colorectal cancer sample. HERGAST demonstrated unique advantages over the other methods. It provided a smoother and more refined delineation of tumor stroma regions, distinguishing them from tumor regions, unlike PCA where stroma regions tended to intermingle with tumors (Figure 2d). This finer resolution is crucial for accurately mapping the tumor microenvironment, enabling better understanding of tumor-stroma interactions for future cancer research^14^. STAGATE(DIC) still suffered from over-smoothing, splitting normal colon mucosal tissue into different clusters (right panel of Figure 2c, box ➀) and disrupting the continuity of tumor areas (right panel of Figure 2c, box ➁, ➂ and ➃). Within tumor regions, HERGAST identified a distinct gene expression profile area.

Highly expressed genes in this region were normalized and merged into a metagene, showing an evident spatial distribution pattern (Figure 2e). Enrichment analysis revealed significant upregulation of immune-related pathways, indicating enhanced anti-tumor immune response and active infiltration of immune cells within the tumor microenvironment^15^. HERGAST also identified a unique cluster of SPP1+ macrophages surrounding calcified areas (Figure 2f, Figure S3 and S4b-h), which were validated as phagocytes through correlation with HE pathological images and confirmation by pathologists. SPP1+ macrophages play a crucial role in the tumor microenvironment and are significant for patient prognosis ^16-18^. In this dataset, SPP1+ macrophages are distributed around tumor cells, forming clusters, strands, or encircling gland-like structures that resemble epithelial cells, posing identification challenges (Figure S4b-g). PCA and STAGATE(DIC) failed to correctly identify these cells. PCA misclassified them as tumor stroma cells, while STAGATE(DIC) grouped them with adjacent cells (Figure S4a, j). In contrast, HERGAST accurately identified these macrophages within the tumor, underscoring its advantage in deciphering complex spatial organization and revealing heterogeneity within tumor regions (Figure S4i). HERGAST’s precise identification of macrophages and immune cells is crucial for understanding the immune landscape of colorectal cancer and informing immunotherapy development ^19^. Similar results were observed on three additional Visium HD datasets (human lung cancer, mouse brain, and mouse small intestine), where HERGAST exhibited more reasonable spatial domain patterns (Figure S8-10)

### HERGAST enhanced critical molecular signature in breast cancer

Next, we analyzed Xenium, another large-scale spatial transcriptomics technology, using a breast cancer tissue section. The spatial clustering results of HERGAST perfectly matched manually annotated regions of interest, successfully identifying and separating invasive cancer, ductal carcinoma in situ (DCIS) #1, and DCIS #2. In contrast, PCA cannot distinguish between the two DCIS areas and STAGATE(DIC) results exhibited excessive regional smoothing, providing limited useful information (Figure 2g). Similar patterns were observed in four other cancer slices (Figure S11-14). To validate the enhancement of significant biological signals in our reconstructed gene expression, we examined the expression patterns of key breast cancer genes (ERBB2, ESR1, PGR, and EGFR) in both the original and reconstructed datasets. The enhanced expression clearly revealed a triple-positive region for ERBB2, ESR1, and PGR, consistent with previous reports^3^ (Figure S5). Additionally, an intriguing observation from the enhanced expression data was the distinct spatial distribution of EGFR and ESR1 within the DCIS regions. EGFR-expressing cells were predominantly located around necrotic areas within the DCIS, forming a clear boundary shape, while ESR1-expressing cells exhibited a diffuse distribution in the DCIS (Figure 2h, Figure S6). Notably, there is little overlap in the spatial expression of these two genes in this slice. This understanding of spatial distribution pattern of the molecular signature can provide insights into microenvironmental influences and cellular interactions within DCIS ^20^, which are not easily discernible from the original expression signals (Figure S7).

## Discussion

In summary, we present HERGAST, an approach for spatial clustering and signal amplification in ultra-large-scale and ultra-high-resolution spatial transcriptomics data. Our method addresses scalability challenges through a divide-iterate-conquer strategy. By integrating gene expression similarity and spatial proximity using a heterogeneous graph network, HERGAST captures local and global spatial relationships effectively, mitigating potential over-smoothing issue. Benchmarking against PCA and STAGATE(DIC), HERGAST consistently outperforms, providing superior spatial clustering and refined delineation of tumor stroma regions. It enables precise identification of high immune-response regions and reveals tumor region heterogeneity, offering valuable insights into the tumor microenvironment and potential immunotherapeutic targets. HERGAST’s enhanced gene expression signal reveals unique spatial expression patterns of key genes in high-resolution and high-noise data, providing insights into microenvironmental influences and cellular interactions. Overall, HERGAST represents a significant advancement in analyzing ultra-large-scale and ultra-high-resolution spatial transcriptomics data, enabling the exploration of complex biological systems with unprecedented resolution.

## Methods

### Construction of relational graph

The spatial neighborhood relationship is established by considering the Euclidean distance between the spatial locations of different spots. In simulation and Visium HD data, we set the graph to include the eight nearest neighbors for each spot. For other datasets, spot *i* and spot *j* are connected if the Euclidean distance between spot *i* and spot *j* is less than a pre-defined hyperparameter, *d*. The expression similarity relationship is constructed by considering the similarity of the PCA representation or selected highly variable genes of the gene expression spectra of different spots. Here we use PCA representation for Visium HD data and all genes’ expression profile for Xenium data. We use Euclidean distance of each spot’s input representation to evaluate the similarity. Across all datasets, we connect each spot to the six spots exhibiting the most similar expression patterns.

### Divide: split ultra-large scale ST data into patches

To scale up the relational graph attention network to arbitrarily large spatial transcriptomics datasets, we devise a “Divide-Iterate-Conquer” (DIC) framework especially crafted for spatial transcriptomics data analysis. We first split the spatial transcriptomics data into multiple patches. The number of patches to split to is a hyper-parameter. If the number of patches is too large, the number of spots in each patch will be too small to contain enough information and cause the training process hard to converge. If the number of patches is too small, the number of spots in each patch will be too large, so the problem comes back to the intractable scale of data. Since most existing methods can handle up to 10,000-20,000 spots per slice, in practice, if the original slice have *n* spots, we suggest splitting the original slice to *m* × *m* patches where 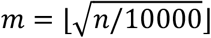 so that on average each patch will have 10,000-20,000 spots. This is also what we adopted in this research.

### Iterate: Iterative training through a relational graph attention network

After splitting original ST slice into several patches, training is iteratively conducted on each patche. As we have discussed before, oversmoothing problem may occur if we cannot carefully design a model that can take both local and global features into account. Here we utilize a heterogeneous graph network that integrates both spatial proximity and gene expression similarity. An intuitive explanation is, a central spot in a small patch can attend to spots at the boundary, which in turn are correlated to neighboring patches through expression similarity (The cells on both sides of the splitting border are mostly the same type of cells). So even if the training process is conducted on small patches, HERGAST model can also implicitly learn some global patterns. This effectively mitigates the issue of over-smoothing that might arise from splitting the data during training.

Through a cross-attention mechanism, our model adaptively learns attention weights for different relationships, automatically weigh the importance of spatial neighbor and expression similarity. The relational graph attention auto-encoder consists of an encoder of two relational graph attention layer and a linear layer decoder.

#### Encoder

The input of the encoder in our architecture is the relational graphs constructed on small patches with *R =* 2 relation types and *N* nodes (spots). The *i*_*th*_ node is represented by a feature vector of the PCA representation or selected highly variable genes expression of ***x***_*i*_. Query and key representations for the *l*_*th*_ encoder layer are computed for each relation type with the help of both query and key kernels, i.e.

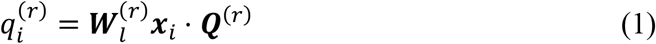

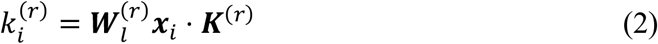

where ***W***_*l*_ is the trainable weight matrix of layer *l*. Then additive attention^21^ is applied to compute attention logits 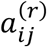:

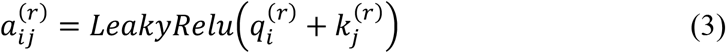

Then the attention coefficients for each relation type are then obtained via the across-relation attention mechanism:

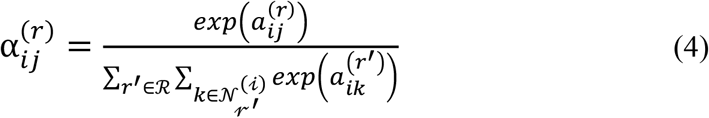

where ℛ denotes the set of relations, i.e. edge types. 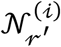 denotes the set of spots connected to node *i* under relation *r*^*′*^. To avoid overfitting, we employed the dropout strategy on the normalized attention coefficients with dropout rate=0.3. That is, 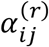 will be set as 0 with a probability of 0.3.

Then spot *i* collectively aggregating information from spots connected to it with the neighborhood aggregation step to get the output of layer *l*. We denote the intermediate representation of 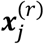 as 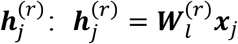. To enhance the discriminative power of the HERGAST layer, we further implement the additive cardinality preservation mechanism^22^:

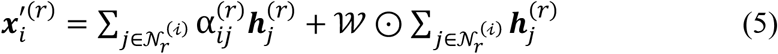

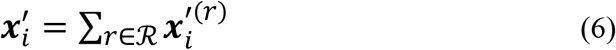

where 𝒲 is a non-zero vector ∈ *R*^*n*^, n is the output dimension of layer *l*, ⊙ denotes the elementwise multiplication.

#### Decoder

The decoder reverses the latent embedding back into the original input representation. The one-layer linear decoder treats the output of the encoder (denoted by ***h***_*i*_) as its input and computes the reconstructed result. Specifically, the decoder computes the reconstructed result as follows:

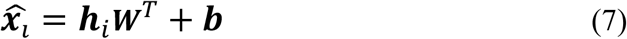

where ***W*** and ***b*** are learnable weight matrix and bias vector. If the input representation is expression of HVG, then a ReLu activation is conducted to get non-negative reconstruction.

#### Loss function

The objective of HERGAST is to minimize the reconstruction loss of the original PCA profiles as follows:

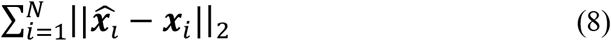

In training process, the model is iteratively trained on each patch. In all the experiments, we used Adam optimizer with learning rate=0.001, weight decay=0.0001 and training epoch=200.

### Conquer: Inference on whole slice using the trained model weights

Since the computation overhead of inference process is much less than training process, inference is conducted on the relational graph conducted based on the entire original dataset. Since the trained model has learned local and global spatial patterns, the output of encoder is considered as the final spot embedding and used to conduct spatial clustering. The output of decoder is considered as the reconstructed amplified gene expression profile.

### Simulation study

For a comprehensive comparison of various methods on simulated data, we utilized the integrated Human Lung Cell Atlas (HLCA) as the single-cell reference dataset. The HLCA consists of over 2.3 million lung single cells with well-established cell type annotations. To simulate different spatial resolutions, we manually designed different spatial patterns with an increasing number of spots and more complex and refined spatial structures. We populated these spatial regions by extracting different cell types from the HLCA and randomly selected one cell type to be diffusely distributed throughout the entire spatial area to mimic widely present cells. To eliminate potential biases from cell type selection, we conducted 10 replicates for each setting. Spatial clustering results were obtained using Leiden algorithm implemented by Scanpy package with resolution=0.01 for all the settings. The performance of the methods was evaluated based on the accordance to the ground truth, with a particular focus on the Adjusted Rand Index (ARI).

### Evaluation metrices

#### Adjusted Rand index (ARI)

The Rand Index computes a similarity measure between two clusterings by considering all pairs of samples and counting pairs that are assigned in the same or different clusters in the predicted and true clusterings. The raw RI score is then “adjusted for chance” into the ARI score using the following scheme:

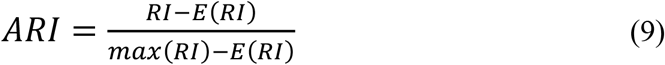

To calculate this value, first calculate the contingency table like that:

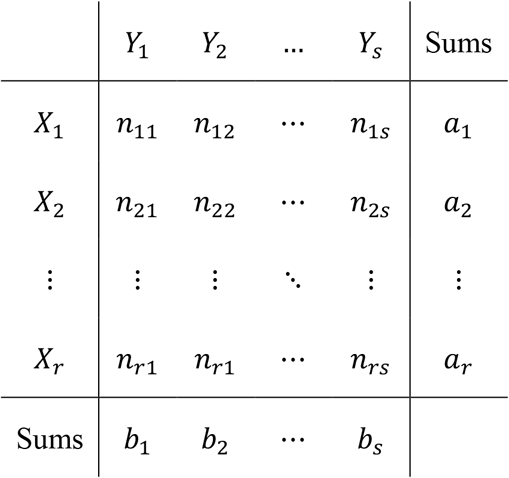

each value in the table represents the number of data point located in both cluster (Y) and true class (X), and then calculate the ARI value through this table:

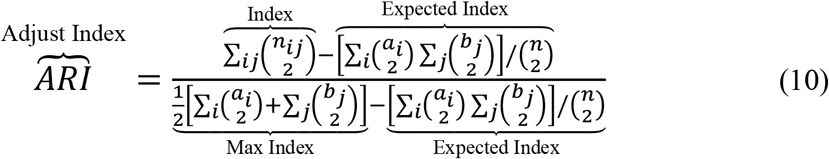

The adjusted Rand index is thus ensured to have a value close to 0.0 for random labeling independently of the number of clusters and samples and exactly 1.0 when the clusterings are identical (up to a permutation). The adjusted Rand index is bounded below by -0.5 for especially discordant clusterings.

### Real data analysis

We applied HERGAST to several ultra-large, super resolution spatial transcriptomics (ST) datasets generated by Visium HD (Human Colorectal Cancer, Human Lung Cancer, Mouse Brain, Mouse Small Intestine) and Xenium (Human Breast Cancer, Human Colorectal Cancer, Human Pancreatic Ductal Adenocarcinoma, Human Lung Cancer, Human Ovarian Cancer). For Visium HD, the 8 µm binned data were used in this study. The data sources and statistics are summarized in Table S1. For Visium HD datasets, we conducted normalization and scaling, and then ran PCA using the Scanpy package^12^. The PCA representation of each spot was selected as the input of HERGAST (Dimension of 200 was used in all Visium HD datasets). For Xenium datasets, the normalized expression of all genes were selected as the input of HERGAST. For all methods, spatial clustering results were obtained using Leiden algorithm implemented by Scanpy package with resolution=0.3 for all the datasets.

## Data availability

All data used in this research are from public database and can be found in Table S1.

## Code availability

Our HERGAST method is openly available from the GitHub repository at https://github.com/GYQ-form/HERGAST.

## Authors’ contributions

Y.G. performed the main research, analyzed data and wrote the original manuscript, X.Y. and Q.J. investigated and interpretated the analysis outcomes. Z.Y. supervised the research. Y.G., X.Y. and Y.Z. discussed and revised the manuscript.

## Acknowledgements

This study was supported by grants from the National Natural Science Foundation of China (Grant No. 12171318 to Z.Y.), the Shanghai Science and Technology Commission (Grant No. 21ZR1436300, 23XD1401900, and 23DZ2290600 to Z.Y.), the Shanghai Jiao Tong University STAR Grant (Grant No. 20190102 to Z.Y.), the Medical Engineering Cross Fund of Shanghai Jiao Tong University (Grant No. YG2023ZD21 to Z.Y.), and Yu Lab. Some parts of computations in this paper were run on the Siyuan cluster supported by the Center for High Performance Computing at Shanghai Jiao Tong University.

