## Supplementary Information for "Unveiling Fine-scale Spatial Structures and Amplifying Gene Expression Signals in Ultra-Large ST slices with HERGAST"

### Supplementary Tables

**Table S1. Detailed description of the datasets used in the paper**

| Datasets | Platform | Data size | Source |
|---|---|---|---|
| Human Colorectal Cancer | Visium HD | 545,913 bins 18,085 genes | https://www.10xgenomics.com/datasets/visium-hd-cytassist-gene-expression-libraries-of-human-crc |
| Human Lung Cancer | Visium HD | 605,471 bins 18,085 genes | https://www.10xgenomics.com/datasets/visium-hd-cytassist-gene-expression-libraries-of-human-lung-cancer-if |
| Mouse Brain | Visium HD | 393,543 bins 19,059 genes | https://www.10xgenomics.com/datasets/visium-hd-cytassist-gene-expression-libraries-of-mouse-brain-he |
| Mouse Small Intestine | Visium HD | 351,817 bins 19,059 genes | https://www.10xgenomics.com/datasets/visium-hd-cytassist-gene-expression-libraries-of-mouse-intestine |
| Human Breast Cancer | Xenium | 167,780 cells 280 genes | https://www.10xgenomics.com/products/xenium-in-situ/preview-dataset-human-breast |
| Human Ovarian Cancer | Xenium | 247,636 cells 380 genes | https://www.10xgenomics.com/datasets/ffpe-human-ovarian-cancer-data-with-human-immuno-oncology-profiling-panel-and-custom-add-on-1-standard |
| Human Lung Cancer | Xenium | 161,000 cells 380 genes | https://www.10xgenomics.com/datasets/ffpe-human-lung-cancer-data-with-human-immuno-oncology-profiling-panel-and-custom-add-on-1-standard |
| Human Pancreatic Ductal Adenocarcinoma | Xenium | 235,099 cells 380 genes | https://www.10xgenomics.com/datasets/ffpe-human-ductal-adenocarcinoma-data-with-human-immuno-oncology-profiling-panel-1-standard |
| Human Colorectal Cancer | Xenium | 388,175 cells 380 genes | https://www.10xgenomics.com/datasets/ffpe-human-colorectal-cancer-data-with-human-immuno-oncology-profiling-panel-and-custom-add-on-1-standard |

### Supplementary Figures

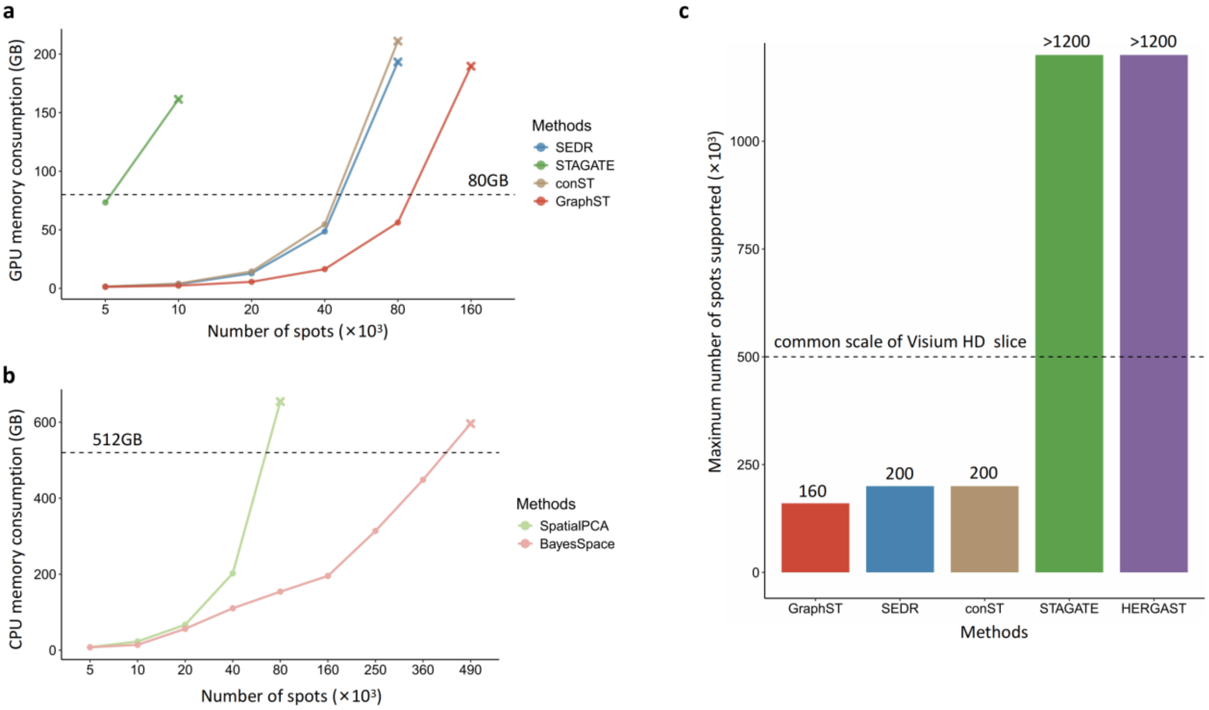


**Figure S1. Scalability comparison on simulated data. a,** Maximum GPU memory consumption of different GPU-based methods in different data scales. Dashed line indicated the maximum cuda memory of GPU in our experiments (NVIDIA A100-SXM4-80GB). Since the last experiment results of each method are out-of-memory and the record cannot be measured, we estimated the approximate required cuda memory based on the cuda memory allocation information obtained during the experiment. **b,** Maximum memory consumption of various CPU-based methods at different data scales. The dashed line represents the memory usage constraint set at 512G. The results of the last experiment for each method exceed the memory limit and cannot be measured, thus are estimated. It is worth noting that while BayesSpace can scale up to 360K spots, it is still not suitable for real Visium HD slices. **c,** Maximum number of supported spots for each method when adopting the DIC strategy. We imposed a maximum memory usage constraint as 512G to guarantee methods' availability. Dashed line indicated the common scale of a Visium HD slice (about 500,000 spots). ">120" means STAGATE(DIC) and HERGAST still not exceeding memory limit when scaling up to 1200,000 spots, which is far more than practical usage.

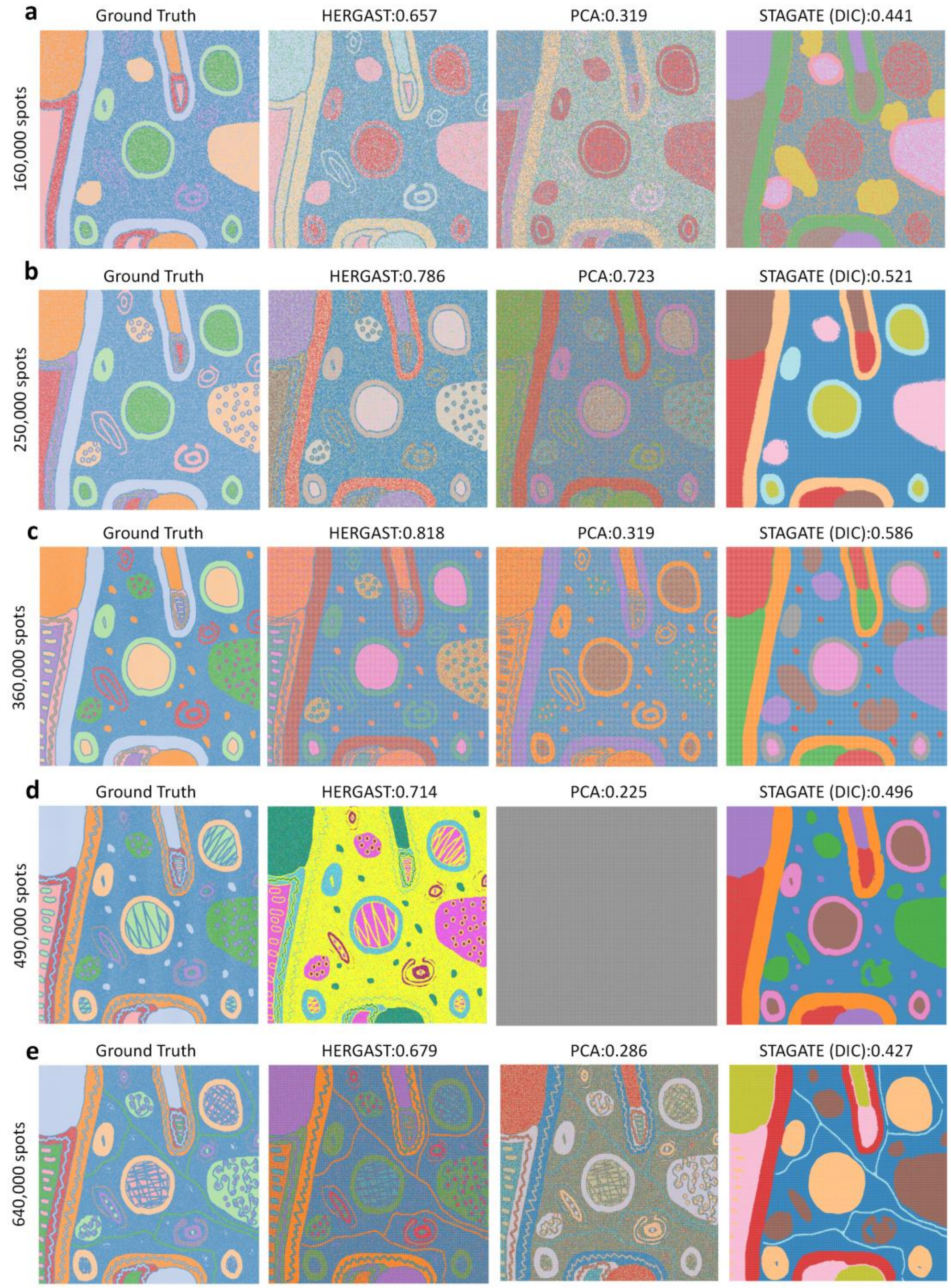


**Figure S2. Simulation results of last replication in each setting.** Each row displays the results of different settings: **a,** 160,000 spots. **b,** 250,000 spots. **c,** 360,000 spots. **d,** 490,000 spots. **e,** 640,000 spots. The first column is the simulated spatial pattern and other three columns are the spatial clustering results of each method. Gray images indicate that there are too many clusters (>100) to display in the panel.

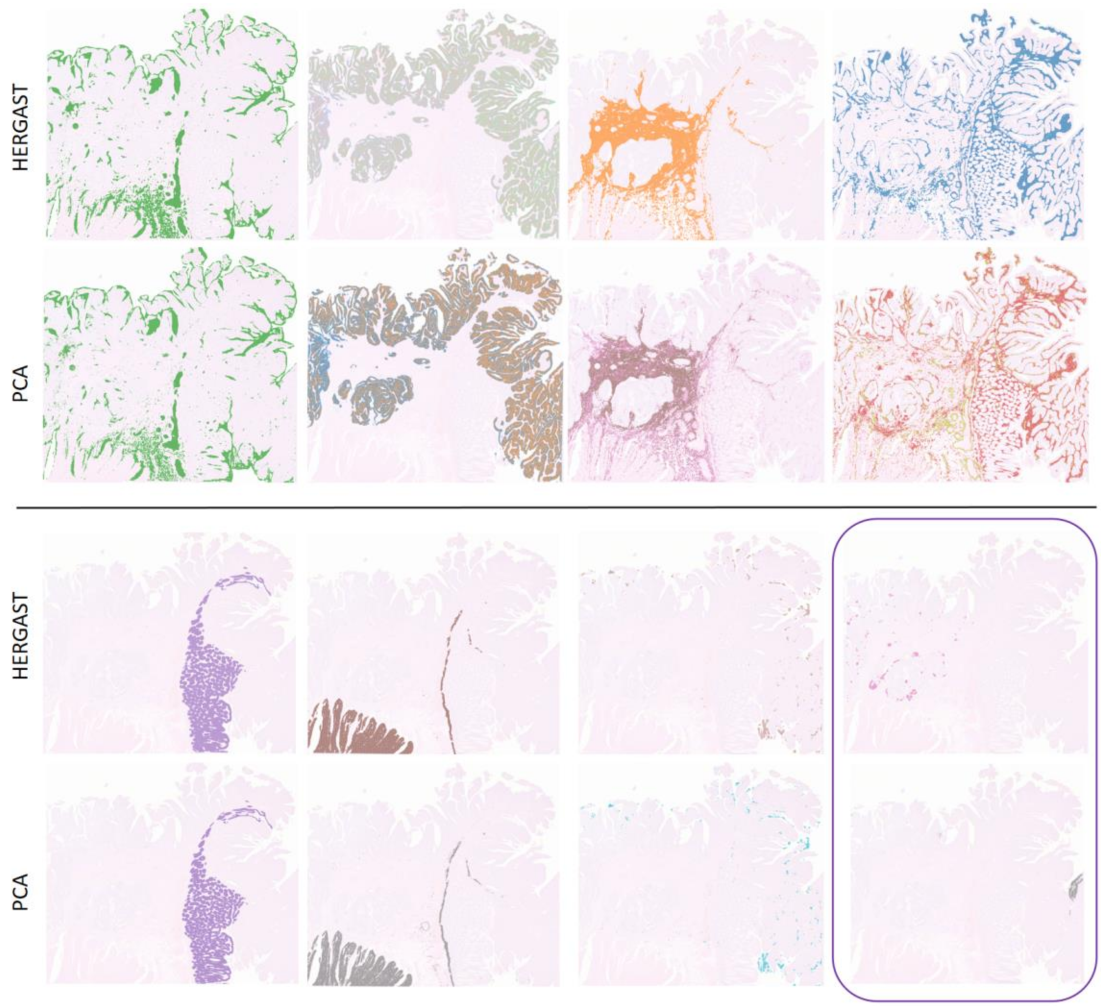


**Figure S3. Match spatial domain between HERGAST and PCA in Visium HD human CRC data.** Each of the first 7 columns is a matching comparison of the spatial clusters obtained by HERGAST and PCA. The last column is two spatial clusters uniquely identified by HERGAST.

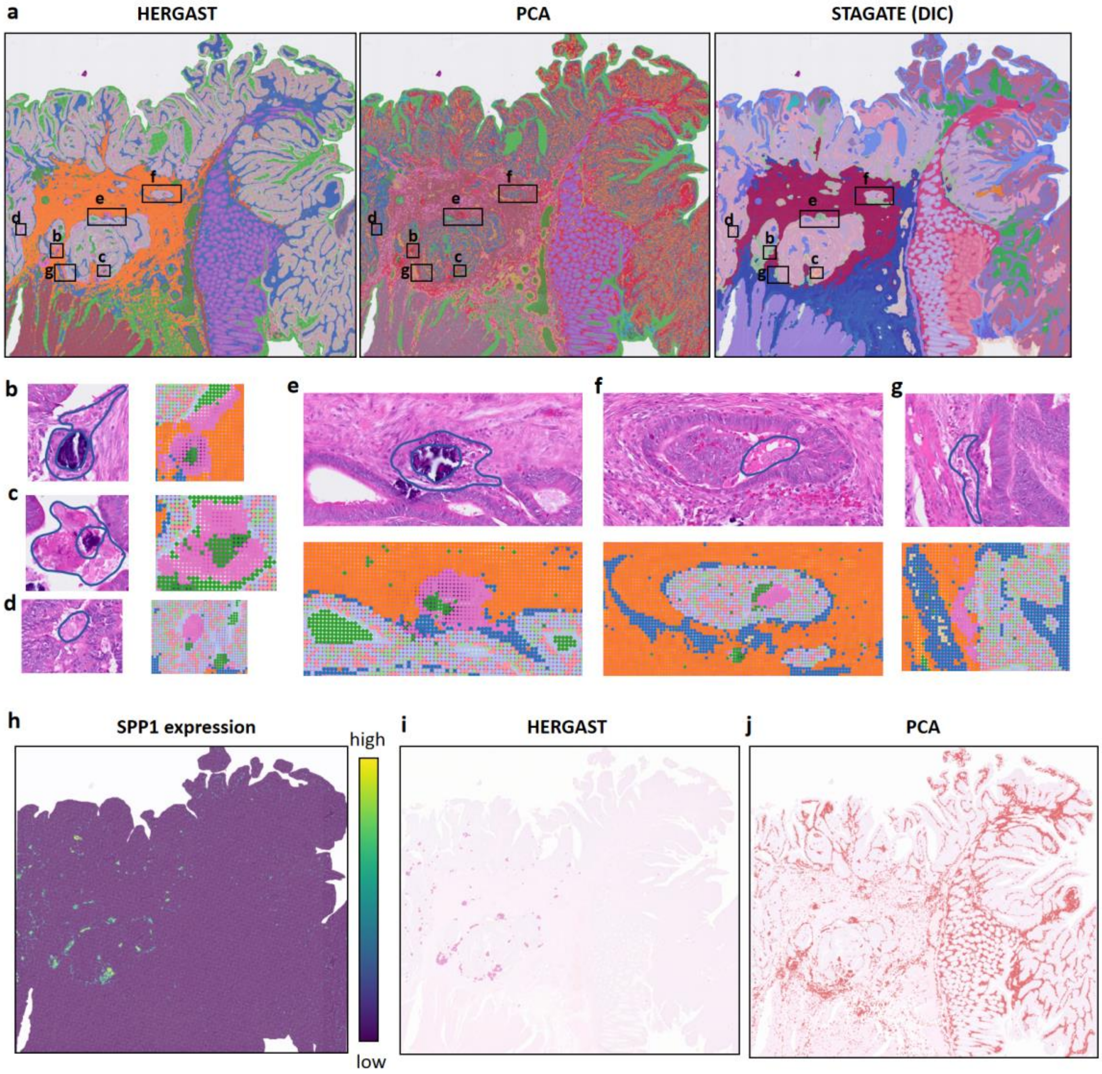


**Figure S4. Unique phagocytes cluster identified by HERGAST. a**, Spatial clustering results of different methods in Visium HD human colorectal cancer slice. **b-g** are the zoom-in views of the corresponding area in **a**, which are selected example of the spatial cluster of phagocytes uniquely identified by HERGAST and the corresponding H&E image. **h**, SPP1 spatial expression plot. **i,** Separate display of the corresponding spatial cluster of HERGAST. **j,** Separate display of the spatial cluster of PCA containing the corresponding area.

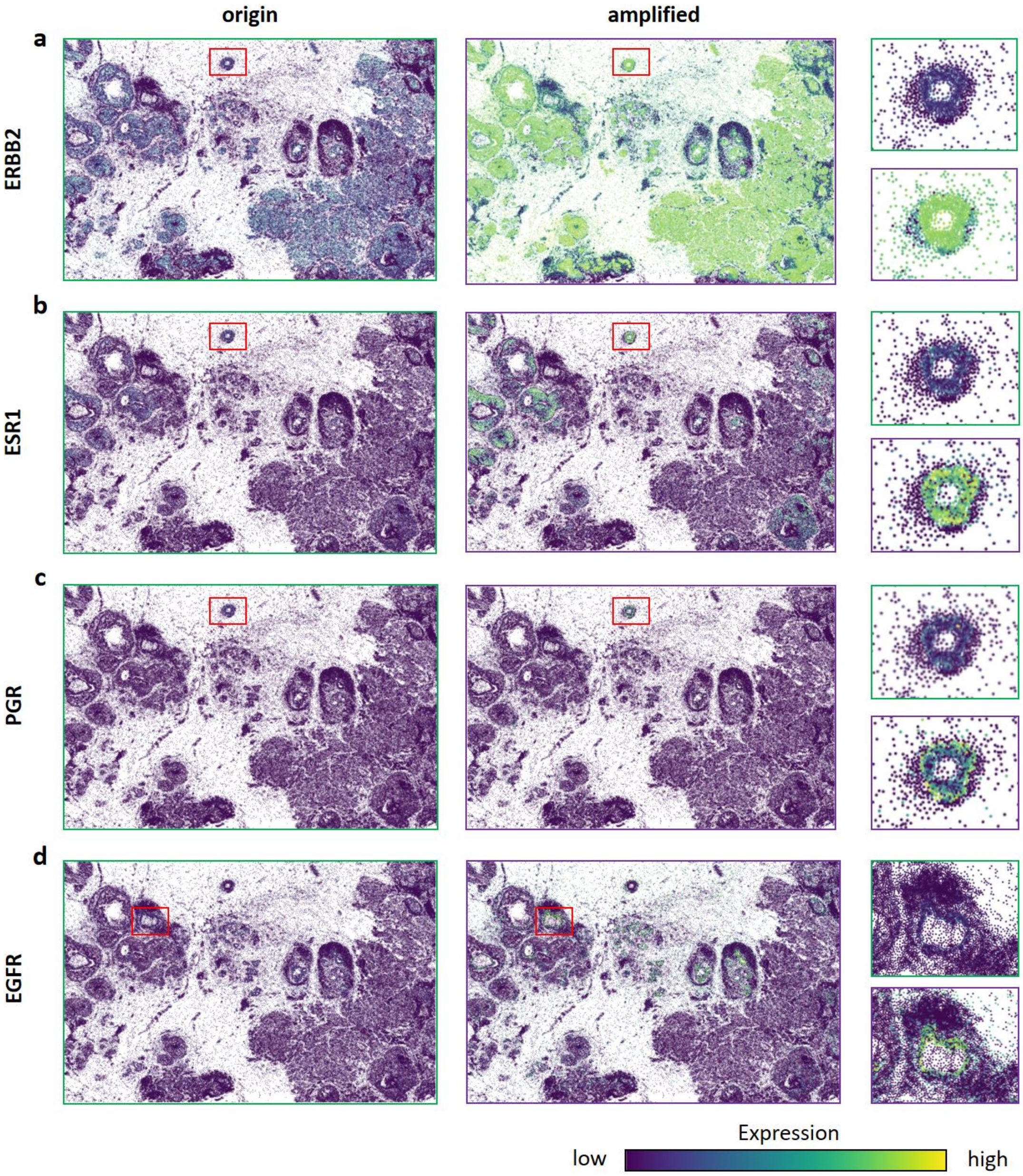


**Figure S5. Spatial expression in original and reconstructed expression profile.** The original and amplified spatial expression of **a,** ERBB2. **b,** ESR1. **c,** PGR and **d,** EGFR. For each gene, the first two columns are the original and amplified expression. The last column represent the zoom-in region indicated in the former slices with top panel displaying original expression and bottom panel displaying amplified expression.

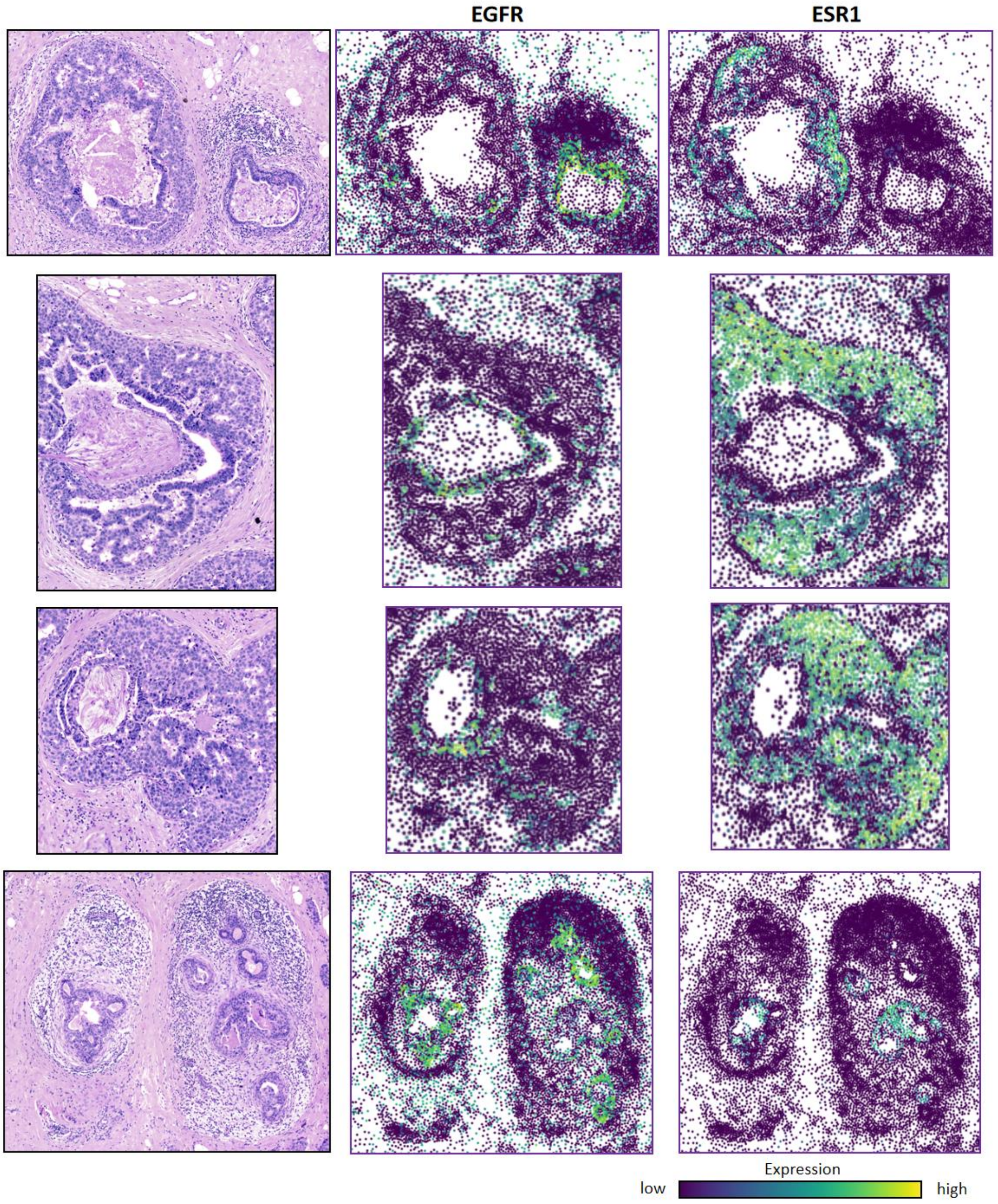


**Figure S6. Amplified spatial expression of EGFR and ESR1 in Xenium breast cancer slice.** The left column is the H&E image of the selected region. Middle and right columns are the amplified spatial expression by HERGAST of EGFR and ESR1 in the corresponding region.

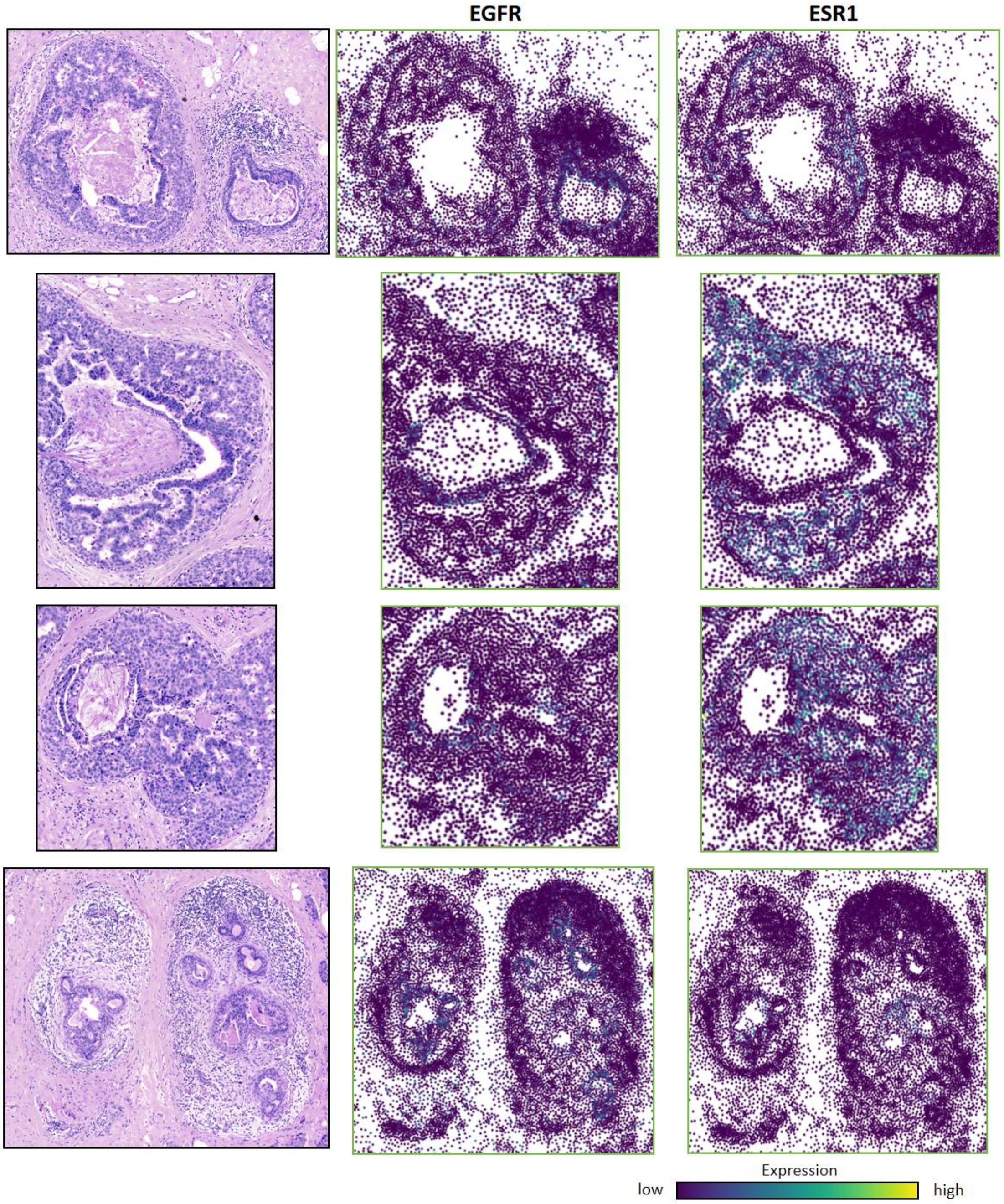


**Figure S7. Original spatial expression of EGFR and ESR1 in Xenium breast cancer slice.** The left column is the H&E image of the selected region. Middle and right columns are the corresponding original spatial expression of EGFR and ESR1 in the corresponding region.

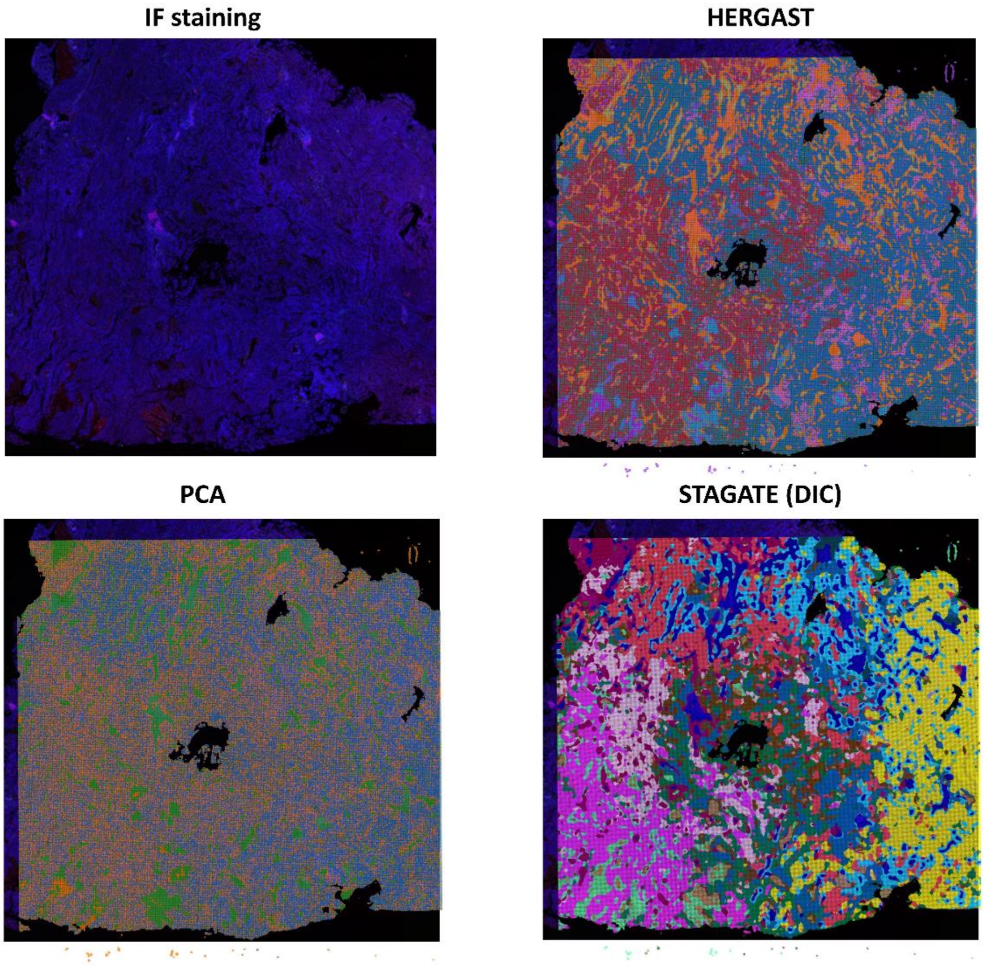


**Figure S8. Spatial clustering results for the Visium HD Human Lung Cancer slice.**

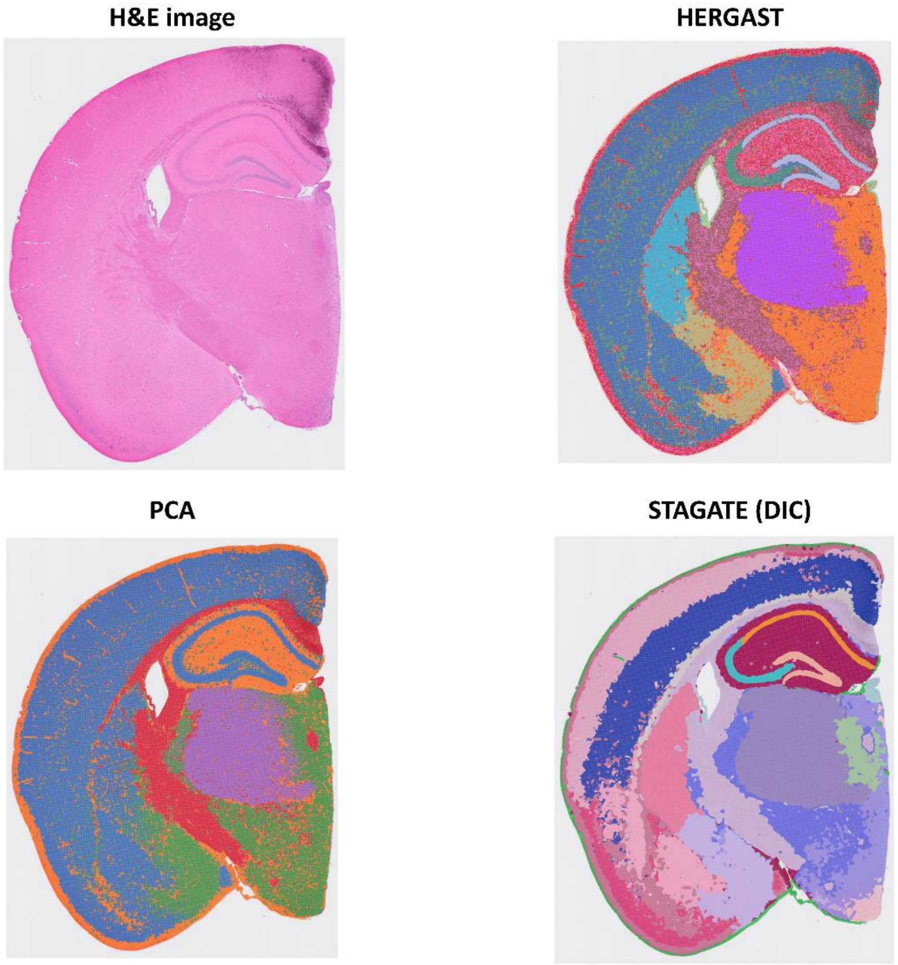


**Figure S9. Spatial clustering results for the Visium HD Mouse Brain slice.**

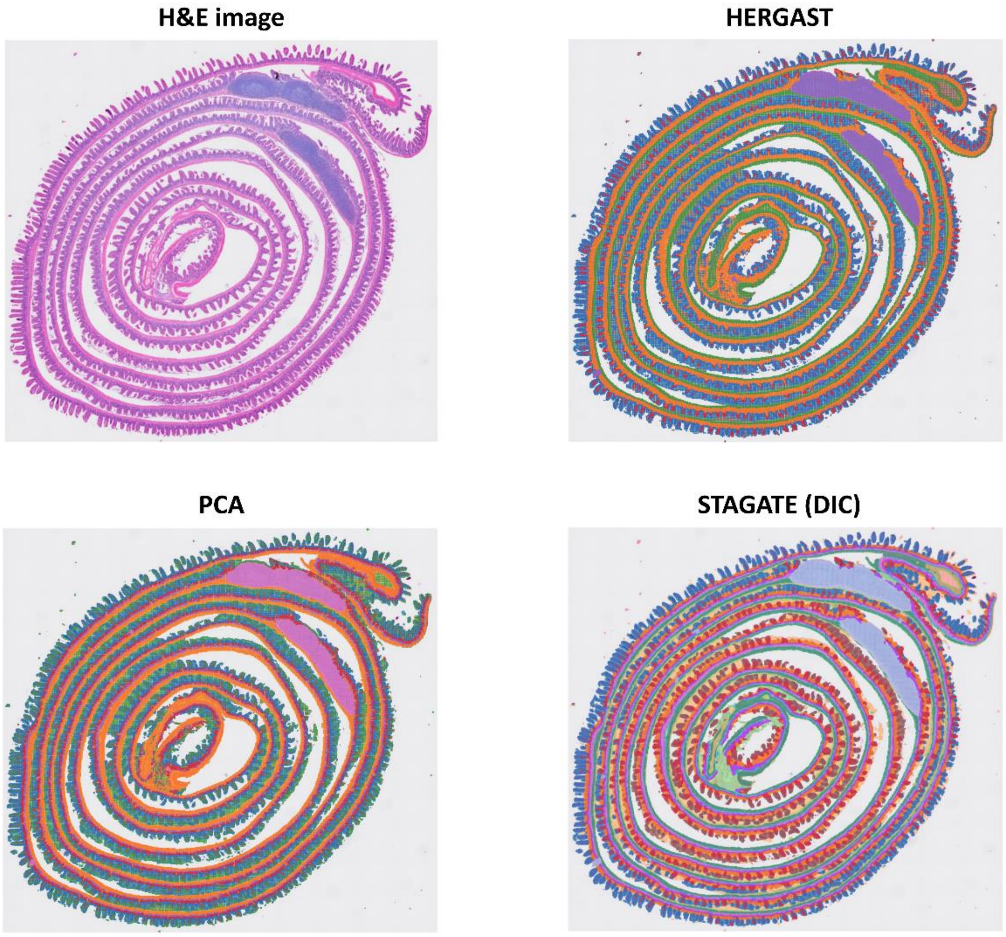


**Figure S10. Spatial clustering results for the Visium HD Mouse Small Intestine slice.**

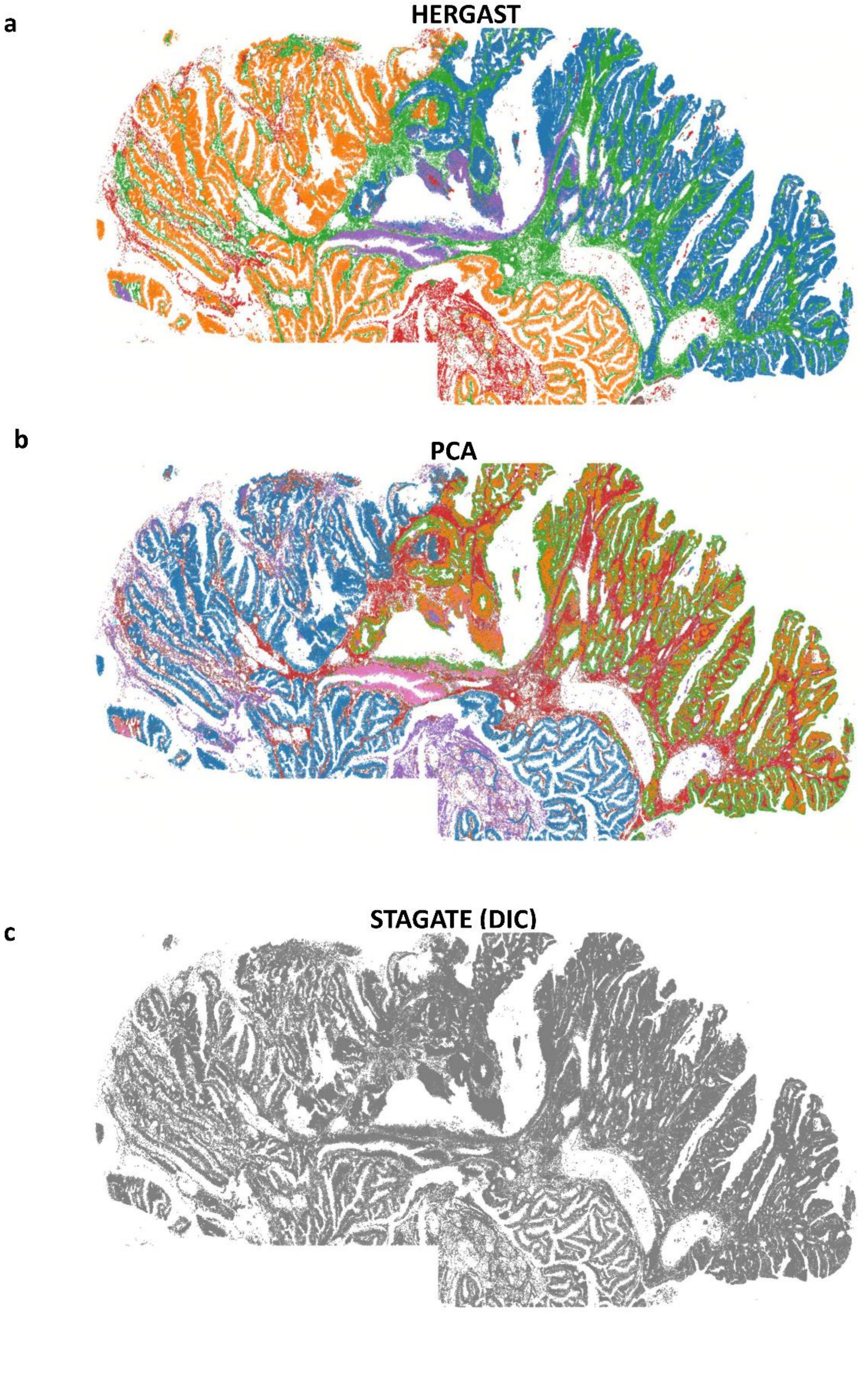


**Figure S11. Spatial clustering results for the Xenium Human Colorectal Cancer slice.** Gray images indicate that there are too many clusters (>100) to display in the panel.

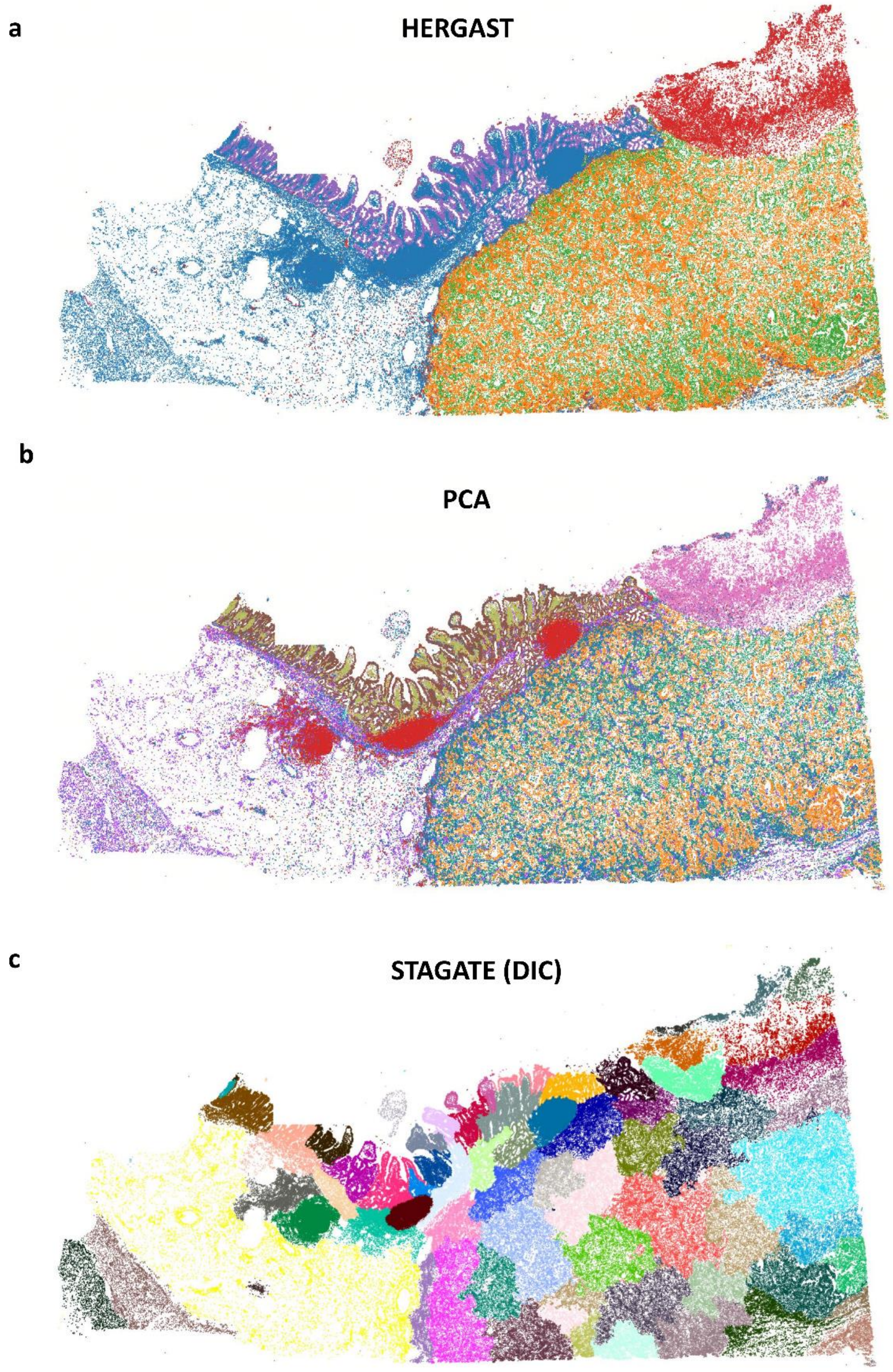


**Figure S12. Spatial clustering results for the Xenium Human Pancreatic Ductal Adenocarcinoma slice.**

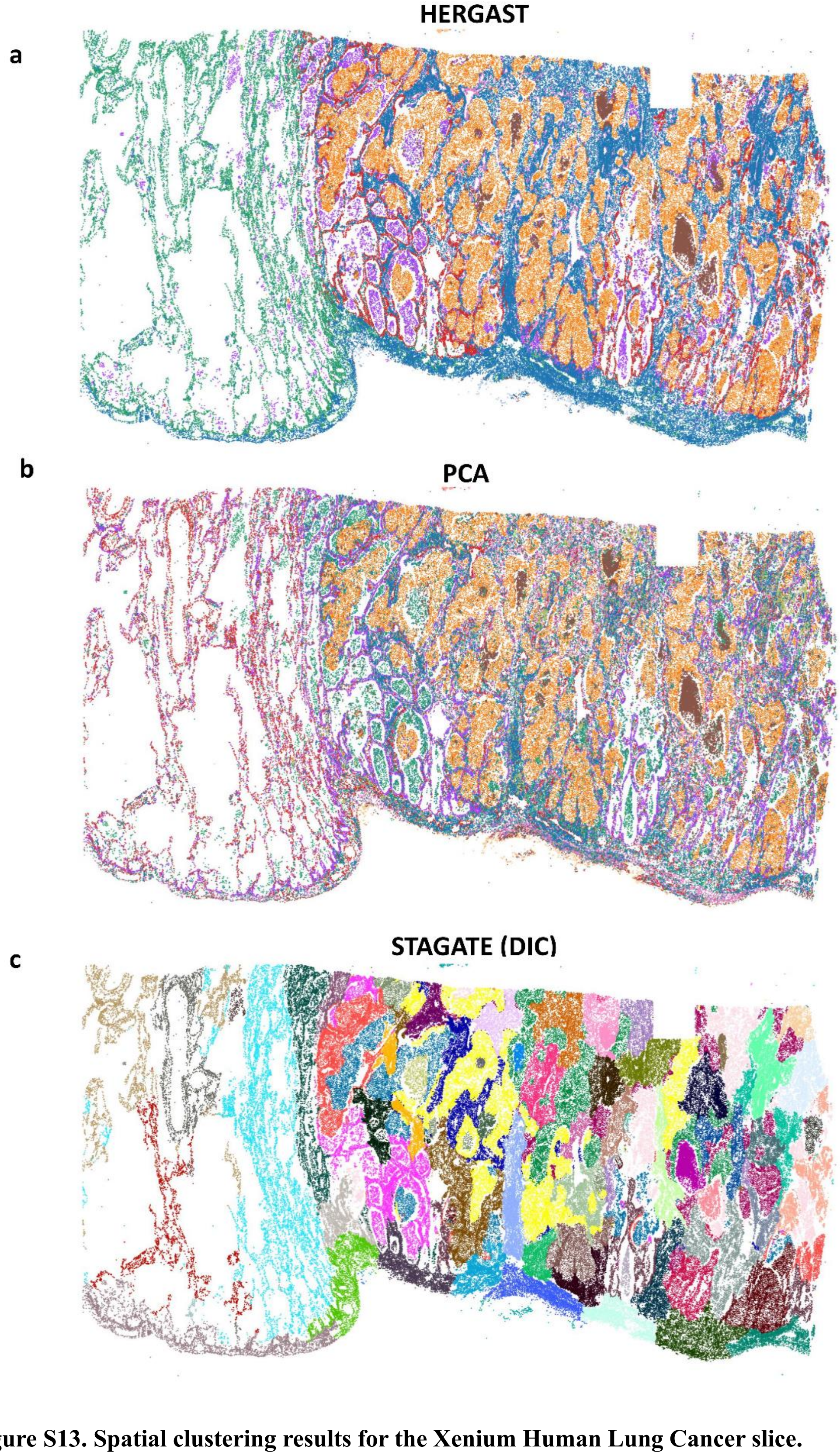


**Figure S13. Spatial clustering results for the Xenium Human Lung Cancer slice.**

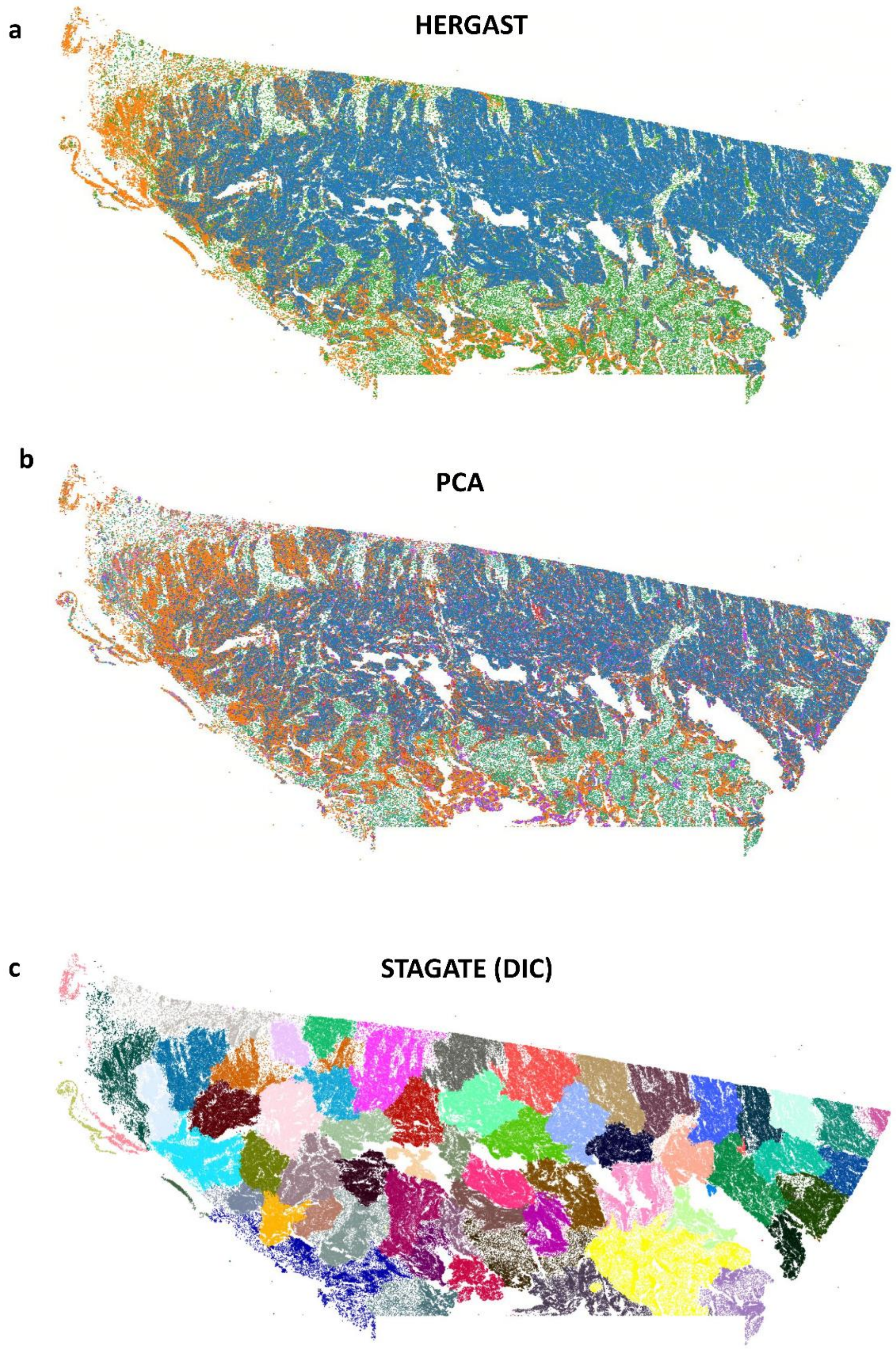


**Figure S14. Spatial clustering results for the Xenium Human Ovarian Cancer slice.**
